# A Mouse Model Of Binge Alcohol Consumption and *Burkholderia* Infection

**DOI:** 10.1101/380683

**Authors:** Victor Jimenez, Ryan Moreno, Erik Settles, Bart J Currie, Paul Keim, Fernando P. Monroy

**Author notes:** Corresponding Author: (FM).

## Abstract

**Background:** Binge drinking, a common form of alcohol consumption, is associated with increased mortality and morbidity; yet, its effects on the immune system’s ability to defend against infectious agents are poorly understood. *Burkholderia pseudomallei*, the causative agent of melioidosis can occur in healthy humans, yet binge alcohol use is progressively being recognized as a major risk factor. Although our previous studies demonstrated that binge alcohol exposure results in reduced alveolar macrophage function and increased *Burkholderia* virulence *in vitro*, no experimental studies have investigated the outcomes of binge alcohol on *Burkholderia* spp. infection *in vivo*.

**Principal Findings:** We used the close genetic relatives of *B. pseudomallei, B. thailandensis* E264 and *B. vietnamiensis*, as useful BSL-2 model systems. Eight-week-old female C57BL/6 mice were administered alcohol comparable to human binge drinking episodes (4.4 g/kg) or PBS intraperitoneally 30 min before a non-lethal intranasal infection. In an initial *B. thailandensis* infection (3 x 10^5^), bacteria accumulated in the lungs and disseminated to the spleen in alcohol administered mice only, compared with PBS treated mice at 24 h post-infection (PI). The greatest bacterial load occurred with *B. vietnamiensis* (1 x 10^6^) in lungs, spleen, and brain tissue by 72 h PI. Pulmonary cytokine expression (TNF-α, GM-CSF) decreased, while splenic cytokine (IL-10) increased in binge drunk mice. Increased lung and brain permeability was observed as early as 2 h post alcohol administration *in vivo.* Trans-epithelial electrical resistance (TEER) was significantly decreased, while intracellular invasion of non-phagocytic cells increased with 0.2% v/v alcohol exposure *in vitro*.

**Conclusions:** Our results indicate that a single binge alcohol dose suppressed innate immune functions and increased the ability of less virulent *Burkholderia* strains to disseminate through increased barrier permeability and intracellular invasion of non-phagocytic cells.

**Author Summary:** *Burkholderia pseudomallei* causes the disease melioidosis, which occurs in most tropical regions across the globe. Exposure rarely evolves to significant disease in the absence of specific comorbidities, such as binge alcohol intoxication. In susceptible hosts, the disease is primarily manifested as pneumonic melioidosis and can be rapidly fatal if untreated. In this study, we utilized *B. thailandensis*, a genetically similar strain to *B. pseudomallei*, and opportunistic *B. vietnamiensis*, a known human pathogen that utilizes similar virulence strategies as *B. pseudomallei* in immunocompromised and cystic fibrosis patients. The study investigates the impact of a single binge alcohol episode on infectivity and immune response *in vivo*. We show that a single binge alcohol episode prior to inhaling *Burkholderia* species increases bacterial spread to the lungs and brain. We also identify alcohol-induced tissue permeability and epithelial cell invasion as modes of action for greater bacterial spread and survival inside the host. Our results support the public health responses being developed in melioidosis-endemic regions that emphasize the nature of binge drinking as a prime concern, especially around potential times of exposure to environmental *B. pseudomallei*.

## Introduction

Binge drinking, and respiratory infections are both significant global health burdens [43]. Patients with alcohol use disorders (AUDs) are more frequently infected with pneumonic pathogens and experience increased morbidity and mortality from these infections [2, 9]. The emerging tropical disease melioidosis is most frequently characterized by pulmonary infection, with pneumonia being the presentation in over half of all cases and a reported mortality rate of up to 53% globally [14]. *Burkholderia pseudomallei* is the causative agent of melioidosis and is a Tier 1 select agent, having been identified as a potential bio-terrorist weapon [50]. The presence of one or more risk factors have been observed in 80% of confirmed melioidosis cases, with nearly 40% of Australian cases having hazardous alcohol use as a risk factor [15]. Similarly, *Streptococcus pneumoniae* is the most common bacterial etiology of community-acquired pneumonia globally and the incidence of pneumococcal infections in individuals with a history of alcohol abuse is higher than the general population [7].

Additionally, hazardous alcohol consumption has been shown to alter the initial host-pathogen interactions during infections caused by *Mycobacterium avium, Escherichia coli, Streptococcus pneumoniae, Klebsiella pneumoniae*, and *Staphylococcus aureus*. [5, 11, 20]. The amount and the pattern of alcohol consumption affect the immune system in an exposure-dependent manner [19]. Most studies indicate acute alcohol consumption is associated with attenuation of the innate inflammatory response expected during infection [41]; whereas, chronic alcoholism produces a predominantly proinflammatory effect that is most often associated with alcohol induced-liver injury [53]. Studies in both human and animal models describe that binge alcohol consumption is characterized by the consumption of 4-6 alcoholic drinks or reaching a minimum blood alcohol concentration (BAC) of 0.08% within a 2 – 3-hour drinking episode [33]. It is unclear specifically how a single binge alcohol intoxication episode alters the lung environment leading to pneumonic infections such as with melioidosis.

In our previous studies, we found binge alcohol conditions alter alveolar macrophage phagocytosis, reactive nitric oxide (RNS) production, and increased intracellular survival of *Burkholderia thailandensis in vitro* [26]. From these findings we concluded that a single exposure of binge alcohol intoxication increased the infectivity of less pathogenic *B. thailandensis* E264 by suppressing the initial host immune response and facilitating a favorable niche for possible bacterial dissemination and survival. However, the *in vivo* effects of binge alcohol consumption on innate immunity during a *Burkholderia* species infection have not been determined. In this study we designed a binge alcohol intoxication mouse model to investigate the effects of a single dose of alcohol on the interaction between less pathogenic *B. thailandensis*, a genetically similar strain to *B. pseudomallei*, and the initial innate immune response to infection. In addition, we utilized opportunistic *B. vietnamiensis* to study the impact of binge alcohol on the infectivity and immune response to a known human pathogen that utilizes similar virulence strategies as *B. pseudomallei* in cystic fibrosis patients [31, 51]. Our results indicate that a single binge alcohol episode can increase *Burkholderia* species infectivity and tissue colonization, while increasing tissue permeability and intracellular invasion of non-phagocytic cells.

## Materials and Methods

### Ethics statement

All use of vertebrate animals at Northern Arizona University was conducted under the American Association for Accreditation of Laboratory Animal Care (AAALAC) regulations and guidelines. Animal care and use were approved in accordance with the Institutional Animal Care and Use Committee (IACUC) according to the policies and procedure of Northern Arizona University (NAU), protocol 16-006. This approval is in accordance with Animal Welfare Assurance A3908-01 from the U.S Department of Health and Human Services.

### Bacterial growth and culture conditions

For each study, frozen stock cultures *(B. thailandensis E264 or B. vietnamiensis Florida, USA strain)* were inoculated into Luria Bertani broth (LB) and incubated overnight at 37°C in an orbital shaker incubator (200 rpm) (New Brunswick C25, Edison, NJ, USA). Bacteria were diluted 1:10 and grown to late-logarithmic phase measured by optical density at OD_600_ absorbance in a spectrophotometer (Eppendorf Bio Photometer AG2233, Hamburg, Germany). Bacteria were collected in 1mL by centrifugation and resuspended in 1mL with pre-warmed Dulbecco’s Phosphate-Buffered Saline (PBS) at a concentration of 1 X 10^5^ or 10^6^ cfu/25μL. Actual numbers of viable bacteria were determined by standard plate counts of the bacterial suspensions on LB agar plates. The Pathogen & Microbiome Institute (PMI), Northern Arizona University, USA, kindly provided *B. vietnamiensis*. All assays were run in triplicate and at least two independent experiments were performed with similar results.

### Animals

Female 8-10 week old C57Bl/6 mice (Jackson Laboratory) with a body weight of 17-21 g were maintained on a standard laboratory chow ad libitum and were housed in a controlled environment with a 12-h light/dark cycle. After receipt, the mice were allowed to acclimate and recover from shipping stress for 5 days in our university laboratory animal facility. These mice were negative for common mouse pathogens during the period of this study.

### Binge alcohol animal model

Binge alcohol intoxication was induced by intraperitoneal (IP) injection of 20% alcohol (4.4 g/kg) in sterile tissue-culture grade water (Sigma Chemical Co., St. Louis, MO) maintained at room temperature. Each mouse was administered a single dose of alcohol that produced a peak BAC of ~256 mg/dL (0.256%). Mice had not been primed previously with alcohol consumption. Control mice received an equal volume of PBS IP. This BAC represents the higher end of the range observed in humans, but it is not particularly rare and has been reported as a common range in human binge drinkers in a number of studies [8]. Notably, mice eliminate alcohol from their systems more rapidly than humans, so producing biologically equivalent effects of alcohol in mice, in order to mimic human binge drinkers, requires a higher dosage in mice. Briefly, viable *B. thailandensis* (10^5^ CFU) or *B. vietnamiensis* (10^6^ CFU) were administered in 25 μl intranasally 30 min after IP injection of alcohol or PBS. Inoculums were administered into each nostril under isoflurane anesthesia. Mice were monitored to observe differences in exploratory and motor control characteristics, in addition to physical health. Mice were subsequently euthanized at 24 and 72 h after the intranasal injection. At these time points, depending on the experimental protocol, aortic blood was taken for cell counts or lung, spleen, and brain tissues were removed and processed for bacterial counts or cytokine measurements. Mice were divided into four groups, and no bacteria was cultured from non-infected mice. At least two independent animal experiments were run with similar results.

### Animal binge alcohol profile and bacteriology of blood and tissues

Blood alcohol concentrations were determined as described in above. Briefly, to determine binge alcohol profile non-infected mice received PBS or alcohol administered as a single dose of 4.4 g/kg of a 20 % (weight/volume) alcohol solution in sterile water by IP injection during the light cycle. Alcohol was injected in mice by using a 27-guage X 0.5-mm (0.4mm X 13mm) needle. All animals were deprived of food and water for 1 h before administration of alcohol but retained free access to food and water post alcohol administration. Blood samples were collected via the tail vein at 30, 60, 360, and 480 min after alcohol administration. Samples were collected in 20 μL heparinized capillary tubes and transferred to 1.5-mL vials that were septum-sealed and stored at 4° C until analyzed.

Blood alcohol concentration measurements were made on blood serum as described in the R-Bio pharm UV-spectrophotometer method protocol (Cat. No. 176 290 035). Analysis was conducted with a UV-Visible Spectrophotometer (Varian Cary 50, Melbourne, Australia).

To quantify bacteria in the blood, blood samples were collected via the tail vein at 2 h PI and were plated directly onto LB agar plates. Plates were incubated overnight at 37° C for quantitative analysis of CFU at 24 and 48 h. Lung, spleen, and brain tissues were aseptically removed at 24 and 72 h PI for quantitative bacterial measurements. Each tissue was weighed in sterile LB and then homogenized with a ceramic bead mix as described by Precellys Lysing Kits manufacturer (Bertin Technologies, Montigny-Le-Bretonneau, France). Samples were then diluted and plated onto LB agar plates. Plates were incubated for 24 and 48 h at 37°C, and CFU counted. Alcohol profile and bacteria assays were run in at least triplicate and at least two independent experiments were performed with similar results.

### GM-CSF, TNF-α, and IL-10 tissue cytokine measurements

Lung and spleen tissue homogenates collected at 24 and 72 h PI were utilized to quantify GM-CSF, TNF-α, and IL-10. Samples were measured using ELISA Ready-SET-Go kits (Affymetrix-eBioscience, San Diego, USA) with procedures supplied by the manufacturer. The minimum detectable levels of GM-CSF, TNF-α, and IL-10 were 4, 8, and <13 pg/mL, respectively. In brief, culture plates were coated with goat anti-mouse GM-CSF, TNF-α, or IL-10 capture antibody and were incubated overnight at 4° C. After the plates were washed, wells were blocked and incubated for 1 h at room temperature. After several washes, respective standards and samples were added to each well, and were incubated overnight at 4° C for maximal sensitivity. After several more washes biotinylated anti-mouse detection antibody was added to each well, and the plate was incubated at room temperature for 1 h. Streptavidin-horseradish peroxidase then was added, and the plate was incubated for 30 min at room temperature. After a final wash, peroxidase substrate TMB solution was added and incubated at room temperature in the dark for 15 min. Adding 3 M sulfuric acid to each well stopped the reaction. Color development in each well was determined spectrophotometrically at 450 nm (Synergy HT, BioTek, Winooski, USA). GM-CSF, TNF-α, or IL-10 results are expressed as pg/mL. Cytokine assays were run in six assay replicates and repeated independently at least twice with similar results.

### Assessment of lung and blood brain barrier (BBB) permeability

To determine the effect of binge alcohol on lung and BBB permeability, mice were administered alcohol as described above. Evans blue dye (Sigma, St. Louis, USA) was administered 30 min post alcohol dose as described by Settles *et al*. (2013). After 1 h, the mice were sacrificed and perfused with 0.9% NaCl saline, and tissues were removed and weighed. Evans blue was extracted by immersing the tissues in a set volume of formamide and extracted Evans blue dye was quantified by measuring the dye absorbance in formamide at 610 nm. Evans blue dye assays were replicated in at least two independent experiments with similar results.

### Measuring transepithelial electrical resistance (TEER) and sodium fluorescein flux in lung epithelial and brain endothelial monolayers

TEER was measured to quantify the effect of a single binge alcohol episode on lung epithelium (EpH4 1424.2, ATCC # CRL-3210) and brain endothelium (bEnd.3, ATCC # CRL-2299). Cell monolayers were grown to confluency on 0.4 micron ThinCert Transwells (Greiner Bio-One, Germany) with DMEM F12 medium (Gibco, Life Technologies) supplemented with 10% fetal bovine serum, 2 mM L-glutamine, 10 mM HEPES, 0.1 mM non-essential amino acids, 1.5 g/l sodium bicarbonate, 50 U/ml penicillin, and 50 mg/ml streptomycin. Cells were incubated at 37°C and 5.5% CO_2_ prior to and after confluency. To characterize the formation of a tight monolayer, TEER measurements were obtained by measuring the overall resistance to the current between electrodes of the insert in a 24-well cell culture plate. The resistance value of a blank culture inserts (reflecting the material, commercial source, and pore size/density used in the experimental case) was subtracted from the total resistance measured, and the resulting value was multiplied by the membrane area to obtain the TEER measurement in Ω cm^2^.

Cell monolayers were incubated in DMEM F12 media supplemented with 0% or 0.2% (v/v) alcohol for 1 and 8 h. Low evaporative cell culture plates and a compensating system were employed as described by Eysseric *et. al*. (1996) [17]. In addition, alcohol- and control non-supplemented media changes were used to ensure consistent alcohol concentrations. Alcohol concentration was selected based on ≥ 90% cell viability utilizing the Trypan blue exclusion cell viability test. Alcohol concentration was also consistent with average mouse BAC. TEER was measured as described previously using a Millicell-ERS Electrical Resistance System (Millipore, Bedford, USA).

To measure monolayer leakiness, cells monolayers grown and treated with DMEM F12 media supplemented with 0% or 0.2% alcohol were rinsed and 0.1 mL of a 10 kDa sodium fluorescein labeled dextran (FITC Dextran) was added to the apical portion of the inserts and 0.6 mL transport buffer (DMEM F12) was added to the basal lateral portion. At 2 h post alcohol administration, 100μL were collected from the basal lateral portion and transferred to a 96 well plate. The 96 well plates were analyzed for fluorescein fluorescence with the Synergy HT micro plate reader. The permeability coefficient of the cells was calculated by subtracting the inverse of the blank coefficient from the inverse of the total permeability coefficient. TEER and FITC-Dextran assays were conducted independently with six assay-replicates and replicated independently twice.

### Binge alcohol and non-phagocytic cells: live Burkholderia intracellular invasion assays

*B. thailandensis* and *B. vietnamiensis* cell invasions with and without alcohol exposure was measured using brain and lung epithelial cells. Briefly, cells were grown as previously described in 24 well cell culture plates. Confluent monolayers were grown in 0% or 0.2% v/v alcohol supplemented DMEM F12 media. *B. thailandensis* or *B. vietnamiensis* were grown overnight in sterile LB media. Prior to co-culturing conditions, the bacteria were diluted to late logarithmic growth, centrifuged, and the pellet was washed twice in fresh non-antibiotic DMEM F12 media. Cell monolayers were then co-cultured with *B. thailandensis* or *B. vietnamiensis* at an MOI of 1:10 for 3 h at 37° C, 5.5% CO_2_ to allow intracellular invasion to occur. After 3 h, extracellular bacteria were removed by washing cells with PBS and replacing culture media supplemented with 250 μg/ml of kanamycin for 1 h. Thereafter, the cell monolayers were incubated (37° C) in media containing 50 μg/ml kanamycin for 1 h for a total of 2 h antibiotic treatment to completely kill any residual extracellular and attached bacteria. Following an additional PBS wash, intracellular bacteria were released after cell monolayers were lysed with PBS containing 0.1% Triton X-100 (total assay incubation time was 5 hours after initial monolayer exposure to bacteria). Viable intracellular bacteria were quantified by plating serial dilutions of the lysate, and average CFU determined. Bacterial intracellular invasion assays were replicated independently at least twice.

### Statistical analysis

The data analysis was completed using Prism 5.0 software (Graph Pad, 5.04, San Diego, CA). Assay replicate independence was determined by a one-way or two-way ANOVA with Bonferroni multiple comparisons, and Student’s *t*-test. Additional statistics were performed using R, and non-parametric, unequal variances. A P value of less than 0.05 was considered significant.

## Results

### C57BL/6 mice binge alcohol concentration profile and quantification of viable bacteria in the blood after infection

To assess the temporal profile of a single binge alcohol episode in female C57BL/6 mice, a dose of 4.4 g/kg alcohol or PBS was administered IP. Blood was sampled and analyzed for blood alcohol concentration (BAC) in mg/dl. A maximum BAC of ~256 mg/dl was generated at 30 min post alcohol administration and was not statistically different at 60 min post alcohol. This dose is consistent with a blood alcohol level that can be achieved during an alcohol binge episode in humans [8]. At 360 min, the BAC declined to 27.6 mg/dl and was not detected (0.0 mg/dl) by 480 min post alcohol (Figure 1A). No alcohol was detected in the non-alcohol administered control mice.

**Figure 1.**
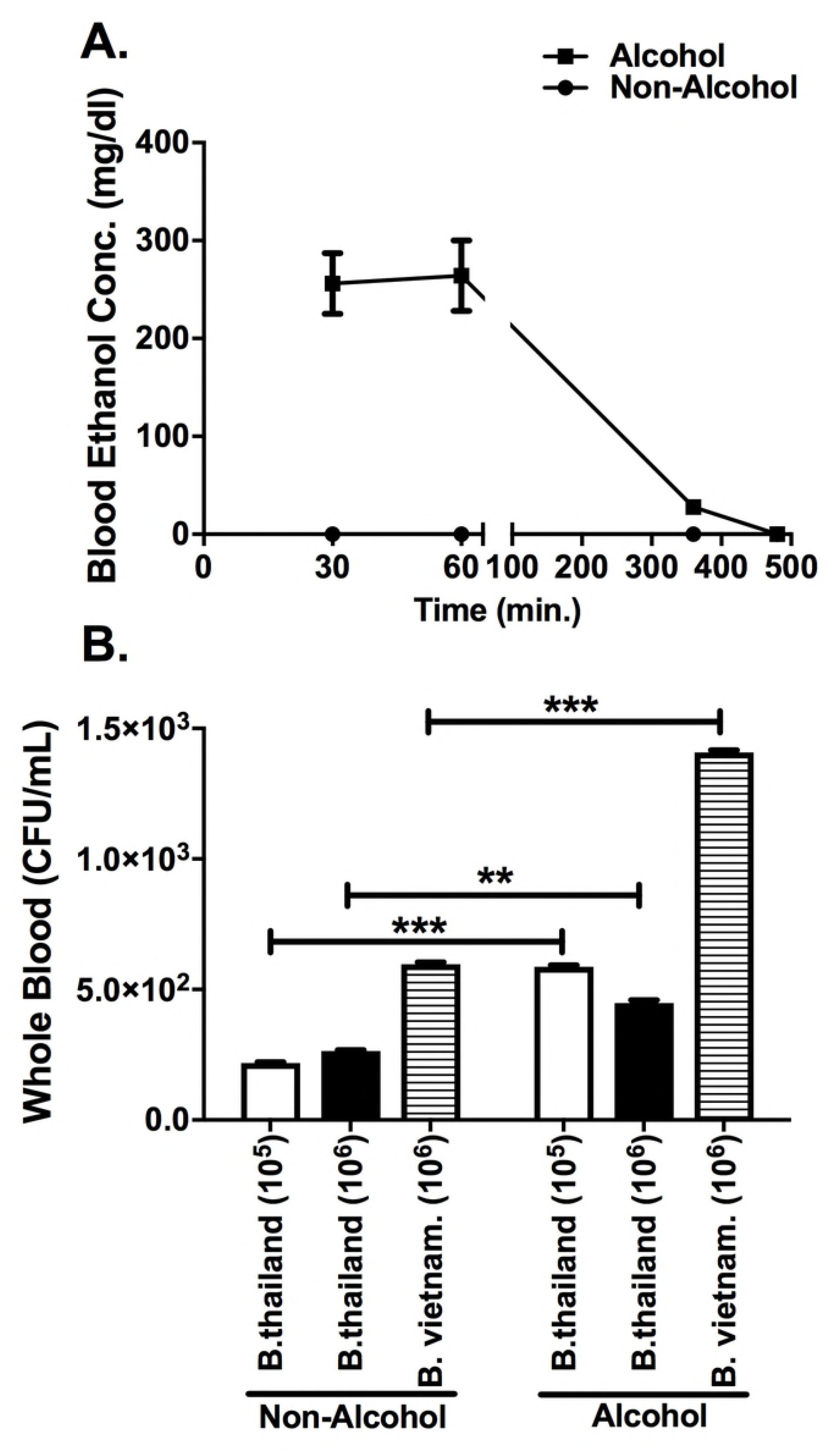
Alcohol and Bacterial Load in Blood. **(A)** Blood alcohol concentration (BAC). C57BL/6 mice were administered alcohol (4.4g/kg) or PBS intraperitoneally (i.p.). Blood was collected for BAC determination at 30, 60, 360, and 480 min post alcohol administration. Trend line represents the average of two independent determinations. n = 3 per determination. **(B)** Mice were administered alcohol or PBS intraperitoneally (i.p.) and 30 min later mice were inoculated intranasally with *B. thailandensis* at doses of (3 x 10^5^), (2 x 10^6^), or *B. vietnamiensis* (2 x 10^6^). Blood was collected at 2 h post infection and *Burkholderia* species were grown on LB media plates to determine colony forming units (CFU). Bars represent the average CFU per treatment with SEM. Horizontal lines and asterisks (*) represent statistical comparison of PBS (Non-Alcohol) control and alcohol treatment determined by Student’s *t*-test, n = 4. **p ≤ 0.01, ***p ≤ 0.001.

We then assessed the ability of *B. thailandensis* and *B. vietnamiensis* to spread to the blood stream with and without alcohol exposure (Figure 1B). Mice were administered a binge alcohol dose 30 min prior to infection, and blood was collected at 2 h PI and immediately spread on LB media plates. Control mice infected with *B. thailandensis* (10^5^ or 10^6^) or *B. vietnamiensis* (10^6^) had significantly less bacteria in the blood compared to infected and alcohol treated mice. On average, 5.8 x 10^2^ CFU’s were collected in the blood from *B. thailandensis* (10^5^) infected mice administered alcohol, compared to a 2-fold decrease in CFU from mice not administered alcohol (*p*= 0.0012). Mice infected with *B. thailandensis* (10^6^) and administered alcohol, developed an average of 4.5 x 10^2^ CFU’s in the blood compared to 2.6 x 10^2^ CFU’s in non-alcohol treated mice (*p*= 0.013). Interestingly, mice infected with *B. vietnamiensis* (10^6^) and administered the same binge alcohol dose developed the greatest bacterial load in the blood of 1.4 x 10^3^ CFU. A 3-fold decrease in whole blood CFU was collected in non-alcohol treated mice infected with *B. vietnamiensis* (10^6^) compared to the binge alcohol treated group (*p*= 0.001) (Figure 1B). Thus, these data suggest that the administration of alcohol increases the dissemination of bacteria shortly after infection.

### Binge alcohol increases bacterial loads in lung, spleen, and brain tissue after Burkholderia intranasal infection

*Burkholderia* infection is reliant on a balance between bacterial dissemination, proliferation, and clearance by host defense mechanisms. A single binge alcohol dose 30 min prior to infection lead to a greater systemic spread and could lead to a greater tissue colonization. Therefore, we investigated tissue burden after 24 and 72 h after infection with and without alcohol administration (Figure 2). A single alcohol dose 30 min prior to infection facilitated a greater bacterial lung, spleen, and brain burden at 24 and 72 h compared to all non-alcohol control groups (Figure 2). More specifically, bacteria were cultured from lung and spleen tissues in binge alcohol treated mice but were not detected (ND) in non-alcohol treated mice infected with *B. thailandensis* (10^5^) 24 h PI (Figure 2A). Conversely, bacterial burden in lung and spleen tissues were detected in non-alcohol treated mice infected with *B. thailandensis* (10^6^) or *B. vietnamiensis* (10^6^) 24 h PI (Figure 2 C, E). Mice infected with *B. thailandensis* (10^6^) or *B. vietnamiensis* (10^6^) and which received the binge alcohol dose presented a 2.5-fold increase in bacterial burden in lung and spleen tissues over non-alcohol treated mice 24 h PI (Figure C, E). No bacteria were cultured from brain tissue in any groups at 24 h PI (Figure 2 A, C, E). Interestingly, bacterial burden in spleen tissue was cleared from non-alcohol and binge alcohol treated mice infected with *B. thailandensis* (10^5^ or 10^6^) (Figure 2 B, D), but detected in spleen tissue of *B. vietnamiensis* infected mice (Figure 2F). Although not statistically significant, greater bacterial burden was measured in lung and brain tissues of mice alcohol treated and *B. thailandensis* infected (10^5^) compared to non-alcohol control mice at 72 h (Figure 2B). Mice that received an increased *B. thailandensis* (10^6^) dose and administered binge alcohol, developed a 4-fold increase in lung and brain bacterial burdens compared to non-alcohol controls (Figure 2D). The greatest bacterial burden (~3 x 10^6^) was localized in lung tissue of mice administered binge alcohol and infected with *B. vietnamiensis* (10^6^) 72 h PI (Figure 2F). Similarly, bacterial burden in lung, spleen and brain tissue of mice administered binge alcohol were significantly greater than non-alcohol control groups of mice infected with *B. vietnamiensis* (10^6^) (Figure 2F). Bacteria were cultured from brain tissue in both groups at 72 h PI (Figure 2 B, D, F). All mice survived by end-point, with mice infected and administered alcohol exhibiting weight loss and lethargy at 72 h PI. These data indicate that the temporal effects of a single binge alcohol episode can increase the dissemination of bacteria in a localized manner.

**Figure 2.**
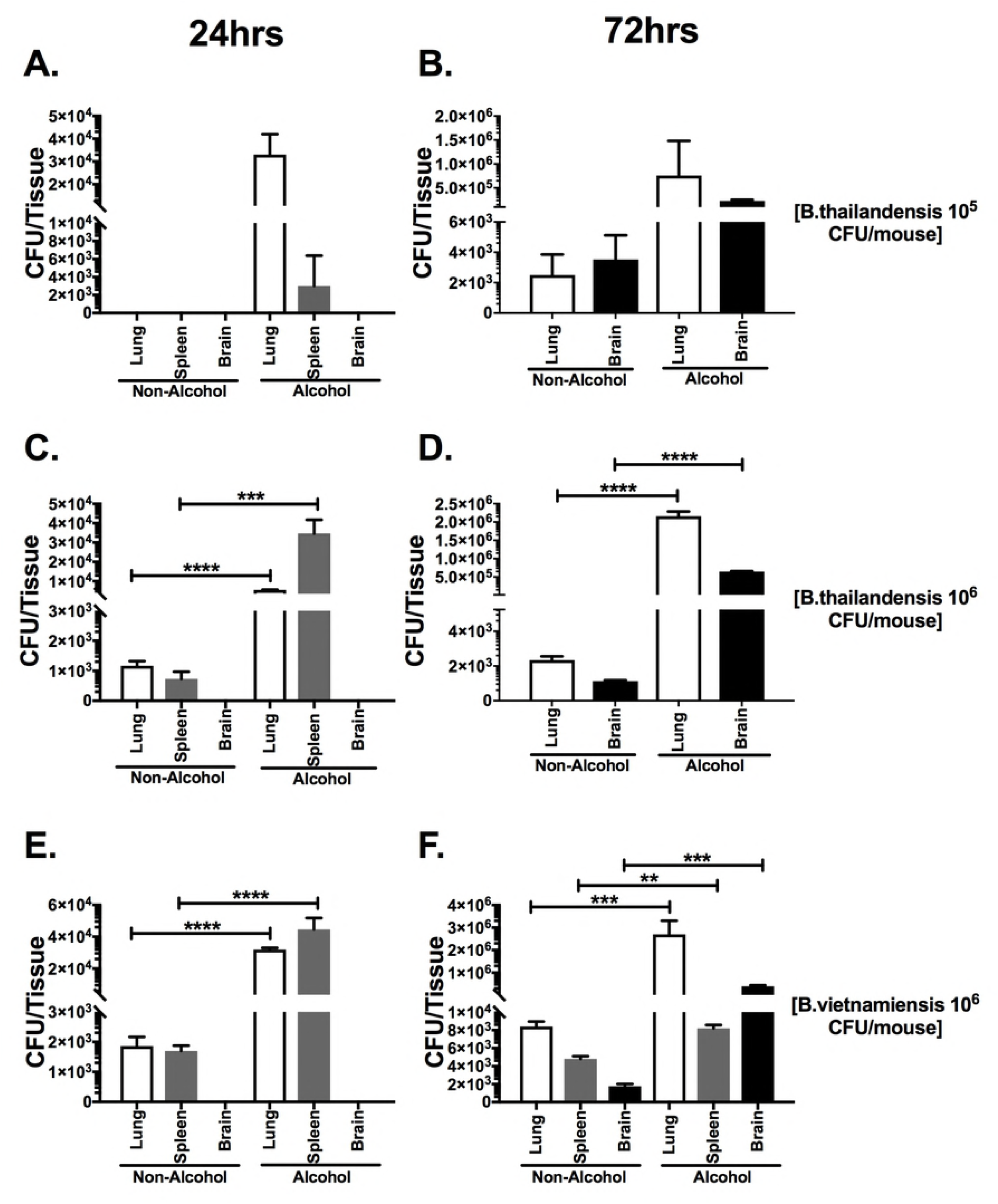
Bacterial load in the lungs, spleen and brain of binge-alcohol mice intranasally infected with different *Burkholderia* species and doses. Mice were administered alcohol (4.4g/kg) or PBS (i.p.). At 30 min. post binge alcohol mice were infected with *B. thailandensis* (3 x 10^5^) **(A-B)**, *B. thailandensis* (2 x 10^6^) **(C-D)**, or *B. vietnamiensis* (2 x 10^6^) **(E-F)**. Tissues were collected 24 h (A, D, C) or 72 h (B, D, F) later and bacterial tissue burden was determined (CFU/tissue). Asterisks (*) represent statistical comparisons between alcohol treatment and (Non-Alcohol) control per tissue type determined by one-way ANOVA. Bars represent average CFU (n=4) with SEM. **, p ≤ 0.01, ***, p ≤ 0.001, ****, p ≤ 0.0001.

### Binge alcohol exposure reduces GM-CSF in the lungs and increases IL-10 in the spleen after Burkholderia infection

The cytokine concentrations were measured at the site of bacterial challenge (the lungs). GM-CSF is produced by alveolar epithelium and binds to specific GM-CSF receptors on the membrane of alveolar macrophages that leads to maturation and differentiation of circulating monocytes. GM-CSF concentrations were decreased in the lungs of mice administered a single binge alcohol dose followed by *B. thailandensis* or *B. vietnamiensis* infections, compared to non-alcohol and infected control mice at 24 and 72 h PI (Figure 3 A, C, E). The greatest decrease in pulmonary GM-CSF was measured in mice administered alcohol and infected with *B. vietnamiensis* (10^6^) at 72 h PI compared to non-alcohol controls (Figure 3E). Decreased concentrations of pulmonary TNF-α were collected in binge alcohol dose mice infected with *B. thailandensis* (10^5^) compared to control mice (Figure 3A). GM-CSF concentrations did not significantly change at 72 h compared to 24 h mice after *B. thailandensis* (10^5^) infection. In mice infected with *B. thailandensis* or *B. vietnamiensis* (10^6^), GM-CSF concentrations were elevated at 72 h above 24 h groups in both non-alcohol and binge alcohol mice (Figure 3 C, E). These data suggest that the alveolar macrophage associated response is dampened in alcohol treated mice.

**Figure 3.**
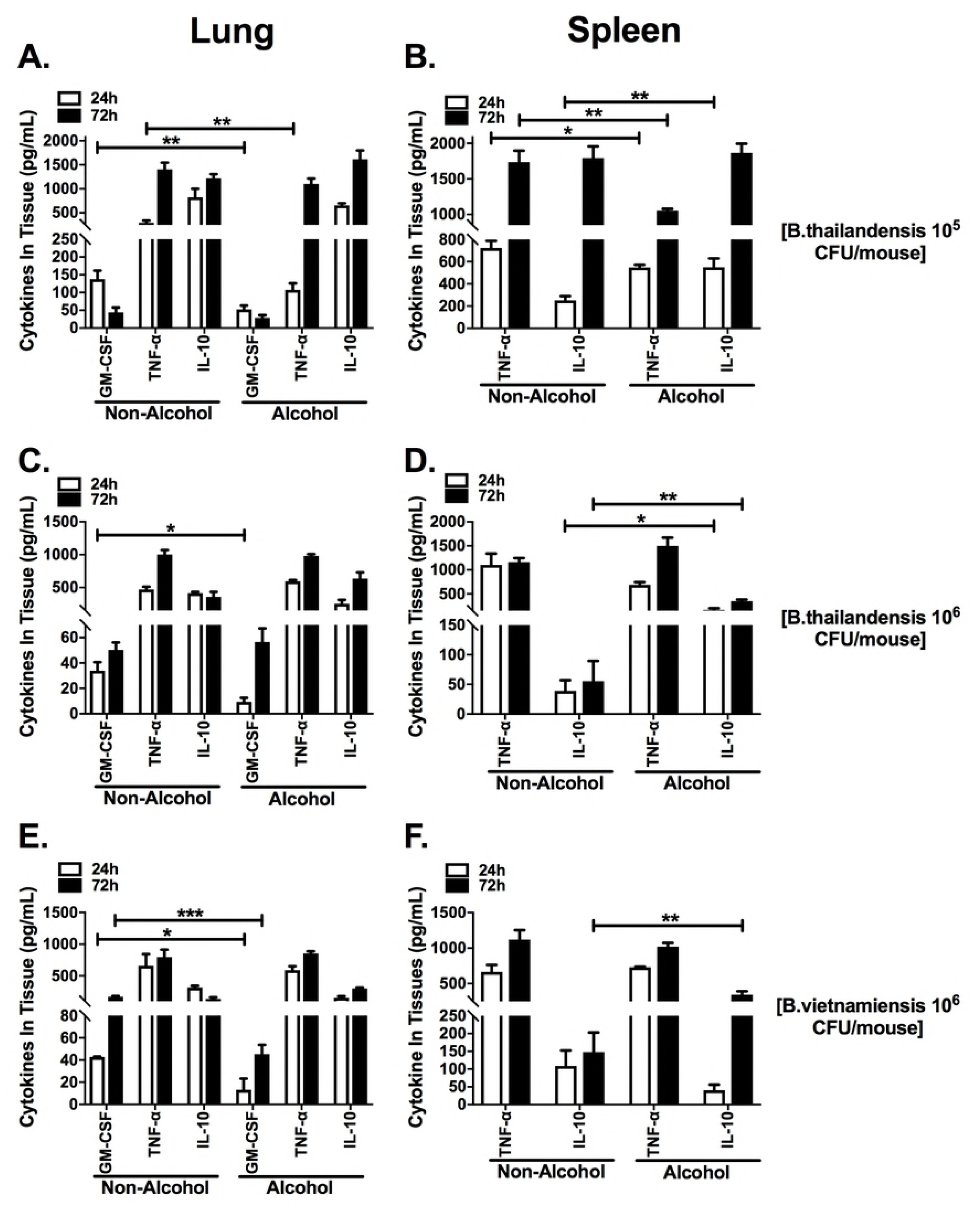
Pro-inflammatory cytokines in lung and spleen of binge alcohol mice intranasally infected with different *Burkholderia* species and doses. Mice were treated as described in Figure 2 and infected with *B. thailandensis* (3 x 10^5^) (**A-B**), B. *thailandensis* (2 x 10^6^) (**C-D**), and *B. vietnamiensis* (2 x 10^6^) (**E-F**). GM-CSF, TNF-α, IL-10 concentrations were measured in lung homogenates (A, C, E). TNF-α and IL-10 concentrations were measured in spleen homogenates (B, D, F), n = 4. Asterisks (*) represent statistical comparisons between alcohol treatment and (Non-Alcohol) control per cytokine determined by two-way ANOVA. Bars represent average concentration (n=?) with SEM indicated. *, p ≤ 0.05, **, p ≤ 0.01, ***, p ≤ 0.001.

Splenic cytokine secretion, a bacterial dissemination site after infection, was also determined in alcohol or PBS administered mice. IL-10 is a modulating cytokine that has been implicated as suppressing the protective immune response, while TNF-α may be immune enhancing in murine models and in humans. Concentrations of IL-10 were elevated in the spleen of mice administered binge alcohol and infected, compared to non-alcohol and infected control mice at 24 and 72 h PI (Figure 3 B, D, F). Interestingly, IL-10 concentrations were elevated only at 24 h PI, while TNF-αwas significantly decreased at 24 and 72 h in mice infected with *B. thailandensis* (10^5^) (Figure 3B). Splenic IL-10 concentrations were significantly elevated in mice administered binge alcohol and infected with *B. thailandensis* or *B. vietnamiensis* (10^6^) at 24 and 72 h or only 72 h PI, respectively. These findings suggest splenic IL-10 may be modulating bacterial clearance in the spleen while dampening the lung immune response in alcohol treated mice.

### A single binge alcohol episode increases lung and brain barrier permeability in vivo

To determine if a single binge alcohol episode could increase lung and brain tissue membrane permeability we used Evans blue dye as an indicator of vascular leakage and tight junction integrity. Mice were administered a single dose of alcohol (4.4g/kg or PBS) 30 min prior to Evans blue dye administration (Figure 4). Mice administered binge alcohol showed increased vascular leakage of Evans blue into lung and brain tissues, however PBS treated mice had reduced infiltration of dye into brain and lung tissue (Figure 4 A, B). Brain tissue from mice administered binge alcohol displayed a 3-fold increase of Evans blue compared to the non-alcohol control mice. Interestingly, more total Evans blue leaked into lung tissue compared to brain tissue of alcohol and non-alcohol treated mice. Mice administered binge alcohol showed greater lung permeability compared to PBS controls, although not statistically significant. In combination, these results suggest that permeability could lead to a greater load of bacteria that is released into the blood.

**Figure 4.**
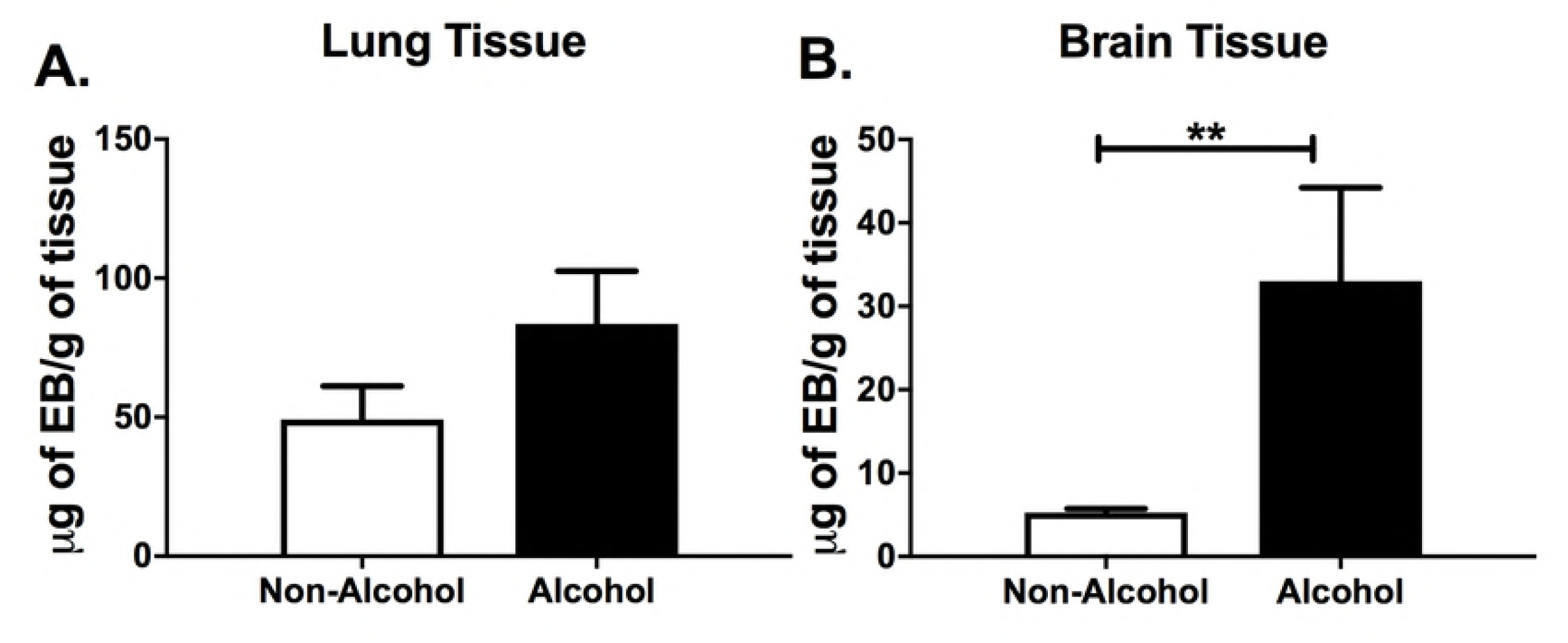
Lung and brain blood barrier permeability in binge alcohol mice. Mice were administered alcohol (4.4 g/kg) or PBS (i.p.). At 30 min. post binge alcohol, mice were injected with Evans blue dye (EB). Mice were sacrificed 2 h post EB administration, perfused, and tissues collected. EB dye was extracted from the **(A)** Lung and **(B)** Brain tissues using formamide. The concentrations of EB per gram of tissue were determined by absorbance at 610nm. Bars represent the average concentration with SEM (n=3). Horizontal line and asterisks (*) represent statistical comparison of PBS (Non-Alcohol) control and alcohol treatment determined by a Student’s t-test, (B) **p ≤ 0.01.

### Binge alcohol reduces transepithelial electrical resistance (TEER) and increases paracellular permeability in lung epithelial and brain endothelial cells

To further investigate the effects of a single binge alcohol episode on tight junction dysfunction and lung and brain tissue permeability, TEER was utilized to measure lung epithelial (Eph4) or brain endothelial (bEnd.3) cell resistance in a transwell system (Figure 5). Both cell types were grown as described in Methods. For the first phase of experiments, membranes with pore diameter size of 0.4 μm were used to support Eph4 cell growth, which represent the lung formed by pulmonary epithelium. The largest average TEER value 2100 ± 25 Ω cm^2^ characterizes the tightest Eph4 cell monolayer. A significant decrease in TEER was observed from the Eph4 monolayer treated with 0.2% (50mM) alcohol v/v at 1 and 8 h post incubation, compared to the no alcohol treated cells (p ≤ 0.0001) (Figure 5A). For the second phase, bEnd.3 brain endothelial cells were also grown on membranes with pore diameter size of 0.4 μm, which represent an acceptable blood brain barrier (BBB) constituent [32]. The largest average TEER value 230 ± 5 Ω cm^2^ characterizes the tightest bEnd.3 cell monolayer. Similarly, an average decrease was recorded from the bEnd.3 monolayer treated with 0.2% (50mM) alcohol v/v at 1 and 8 h compared to the no alcohol treated cells, (p ≤ 0.01) and (p ≤ 0.0001) respectively (Figure 5B).

**Figure 5.**
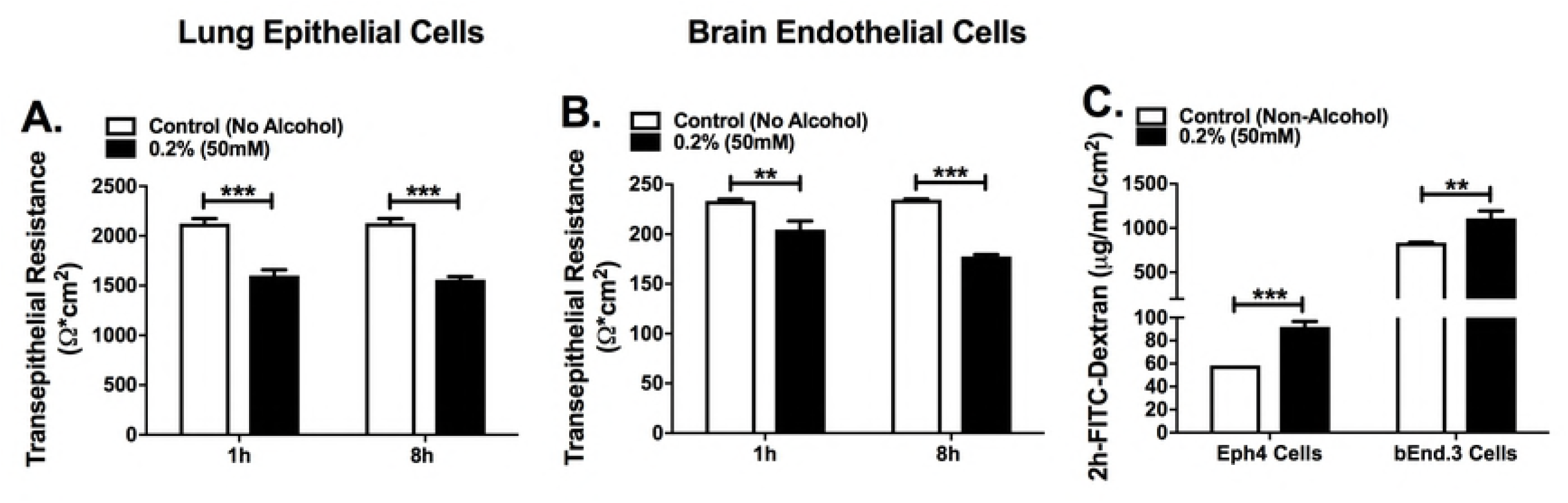
Lung epithelial and brain endothelial cell permeability with and without alcohol treatment. **(A)** Lung epithelial (Eph4) and **(B)** brain endothelial (bEnd.3) cell transepithelial resistance (TEER) was measured in cell monolayers grown in F12 media in 0.4-micron pore diameter membrane inserts. Media was supplemented with 0.0% or 0.2% v/v alcohol and TEER was measured at 1 and 8 h post alcohol administration. In panel (C), the permeability was determined in both cells by adding FITC-Dextran (10 KDa) to the apical side and measured in the baso-lateral side at 2 h post alcohol administration. Bars represent the average TEER across the permeable membrane per treatment with SEM. Horizontal lines and asterisks (*) represent statistical comparison of PBS (Non-Alcohol) control and alcohol treatment determined by Student’s t-test at each time point, (A-C) **p ≤ 0.01, ***p ≤ 0.0001.

Cell monolayers were also characterized by FITC – Dextran permeability. A decrease in TEER value for binge alcohol dose treated cells was associated with an increase in FITC-Dextran paracellular permeability with both lung and brain cells supplemented with 0.2% v/v alcohol compared to non-alcohol controls (*p* ≤ 0.0001) and (*p* ≤ 0.001) respectively (Figure 5C). No cell death was observed in the monolayers treated with the binge alcohol dose. Taken together, these results propose bacterial bi-directional diffusion across paracellular space in mice administered alcohol.

### Binge alcohol increases intracellular invasion of lung epithelial and brain endothelial cells

To further examine the paracellular diffusion results, non-phagocytic lung epithelial and brain endothelial cells were tested for intracellular invasion susceptibility during a binge alcohol dose. Monolayers were formed and co-cultured in with or without alcohol. The results in Fig. 6 show the average number of CFUs, demonstrating viable *B. thailandensis* or *B. vietnamiensis* isolated 3 h after challenge. Although not statistically significant, *B. thailandensis* infected lung epithelial cells treated with binge alcohol resulted in a ~ 2-fold increase in intracellular invasion compared to non-alcohol treated cells. *B. vietnamiensis* invasion of binge alcohol treated lung epithelial cells resulted in an ~ 3.5-fold increase compared to non-alcohol treated cells (Figure 6A). Both *Burkholderia* strains increased effective intracellular invasion and survival when alcohol was present (Fig. 6 A, B). Comparably, a ~3-fold increase in viable *B. thailandensis* was recovered from brain endothelial cells treated with binge alcohol, while a 2-fold increase in intracellular invasion resulted from *B. vietnamiensis* infected brain endothelial cells treated with binge alcohol, compared to non-alcohol treated cells (Figure 6B). Intriguingly, brain endothelial cells were more susceptible to *B. vietnamiensis* intracellular invasion compared to *B. thailandensis* in both alcohol and non-alcohol groups (statistical comparison not shown), with greater bacteria recovered from binge alcohol treated brain endothelial cells (Fig. 6B). These findings suggest a single binge alcohol episode increases bacterial survival and dissemination through an increase in intracellular invasion of non-phagocytic cells.

**Figure 6.**
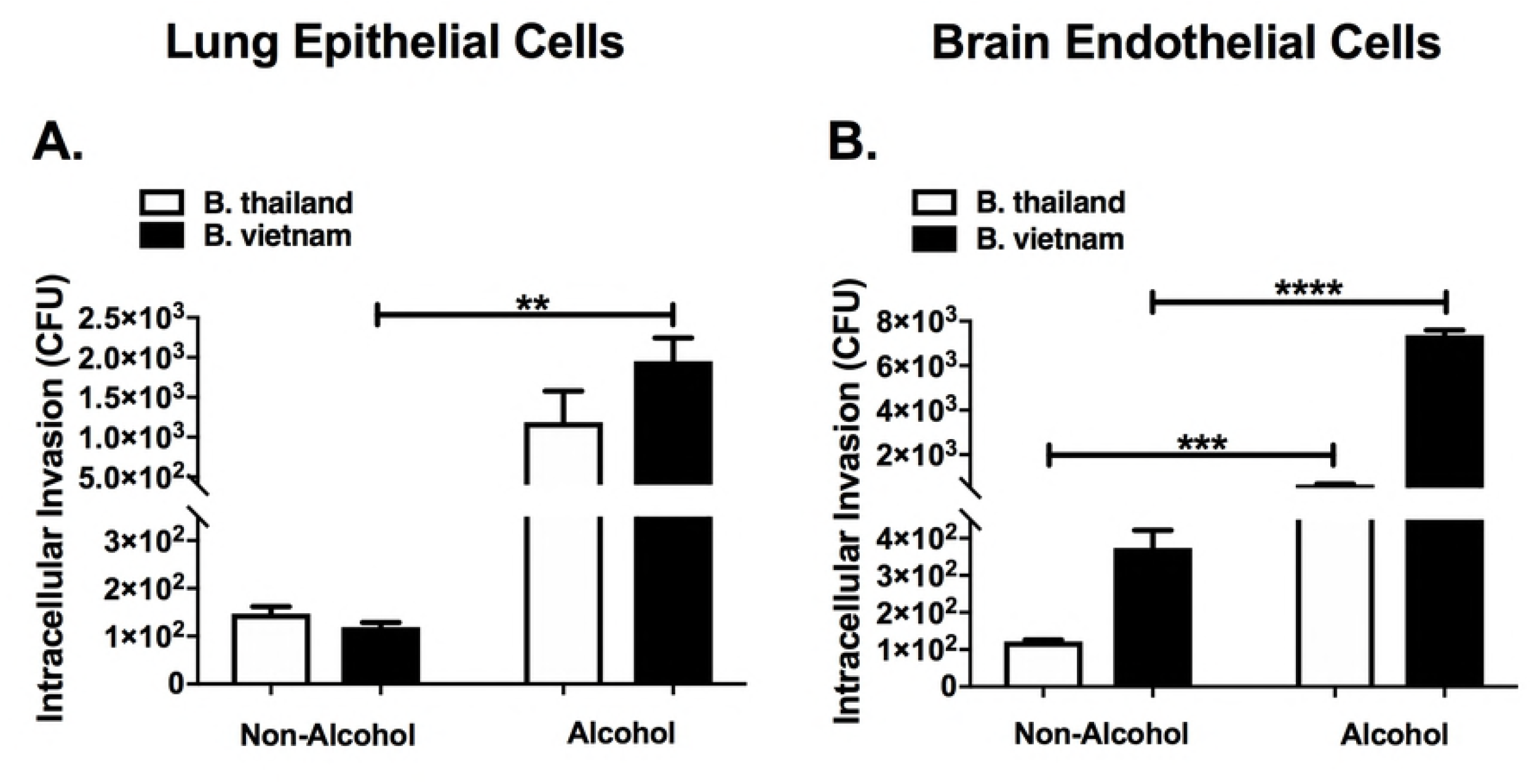
Bacterial invasion and survival in non-phagocytic lung and brain cells with and without alcohol treatment. **(A)** Lung epithelial and **(B)** brain endothelial cells were grown to confluency in F12 cell culture media and co-cultured with *B. thailandensis* or *B. vietnamiensis* (MOI 1:10) for 3 h in media supplemented with 0.0% or 0.2% v/v alcohol. Extracellular bacteria were removed by washes X4 and antibiotic treatment for 2 h. Cells were lysed and viable bacteria recovered. Asterisks (*) represent statistical comparisons between alcohol treatment and (Non-Alcohol) control determined by one-way ANOVA. Bars represent average CFU with SEM. **, p ≤ .01; ***, p ≤ .001; ****, p ≤ 0.0001.

## Discussion

Binge alcohol intoxication has been recognized as a risk factor for infections including pneumonia and other sepsis [38]. Our previous studies have indicated that binge alcohol exposure of alveolar macrophages before or after infection decreased resistance to infection, in part by decreasing inflammatory mediators and phagocytic mechanisms [26]. Melioidosis has been linked to binge alcohol use through epidemiological studies, but the effects of binge alcohol exposure on pathogenic or less-pathogenic *Burkholderia* strains and innate immunity have not been examined *in vivo*. This was the basis for our working hypothesis that the same innate dysfunction observed *in vitro* would occur in our binge alcohol mouse model. In addition, we hypothesized that a lone binge alcohol episode would enhance the virulence of less pathogenic and opportunistic *Burkholderia* spp.

In the present study we used the less-pathogenic *B. thailandensis* E264 and the opportunistic *B. vietnamiensis*, a strain that infects cystic fibrosis patients, as models to study the effects of binge alcohol consumption on infection. We observed an increase in *Burkholderia* species dissemination into the blood as early as 2 h PI, and greater bacterial loads in lung and spleen tissues 24 h PI in mice that had blood alcohol levels equivalent to a binge alcohol episode compared to mice that did not receive alcohol. Interestingly, the binge alcohol dose administered was completely cleared from the blood of mice by 8 h post administration (Figure 1), and yet a clear dysfunction of bacterial clearance from tissues was still dampened or delayed (16 and 64 h after alcohol was cleared).

Sufficient inflammatory infiltrates in non-alcohol treated C57BL/6 mice may explain, in part, the clearance of bacteria from tissues with *B. thailandensis* infections at 10^5^ or 10^6^ CFU. Similarly, other groups using C57BL/6 intranasal mouse models have reported that infiltration of macrophages within the first 3 days of infection may serve to contain *B. thailandensis or B. pseudomallei* for a longer period than in BALB/c mice, allowing the initiation of an adequate immune response (16, 36). The ability of *B. vietnamiensis* and other closely related *B. cenocepacia* species to cause severe infection in patients with cystic fibrosis led to the use of a Florida, USA strain collected from soil [22]. A more complete understanding of the effects of binge alcohol on a known human pathogen would potentially allow for the development of effective preventative strategies for highly virulent *B. pseudomallei*. Mice exposed to alcohol and infected with *B. vietnamiensis* exhibited greater bacterial colonization of the lung and spleen at 24 h PI and retained significant bacterial loads in the spleen at 72 h PI, compared to mice infected with *B. thailandensis* (Fig. 2). Considering the rapid immunological response and tolerance of C57BL/6 mice to *Burkholderia* species infections, these findings reveal that a single binge alcohol episode can increase tissue colonization while suppressing innate immunity in both *B. thailandensis* and *B. vietnamiensis* infections.

To better understand tissue colonization and the modulatory effects of binge alcohol on innate immunity, tissue cytokines were examined. The cytokine, GM-CSF has a dual role in augmenting the accumulation and activation of both neutrophils and macrophages that boosts the infection-fighting ability of host lung defenses [35, 37]. We have observed previously that macrophages are dysfunctional after alcohol treatment *in vitro* [26]. From our study, GM-CSF and modulating cytokine TNF-α were elevated as early at 24 h in lung and spleen tissues in mice infected with *B. thailandensis* or *B. vietnamiensis* at differing doses in the absence of alcohol compared to mice that received alcohol (Figure 3). Moreover, binge alcohol administration consistently reduced GM-CSF with mild effects on TNF-α in the lungs, compared to elevated levels in non-binge drunk mice. These pulmonary results indicate a reduction in activated neutrophils and macrophages [28]. Intriguingly, elevated IL-10 in the spleen of binge drunk mice infected with *B. thailandensis* may provide insight into the modulatory effect of binge alcohol on the general inflammatory cytokine profile of innate immune cells [13]. Although outside the scope of this study, it is plausible that a cytokine “storm” may be induced in the spleen of C57BL/6 mice during prolonged infections. Binge alcohol may directly or indirectly mitigate these detrimental cytokine effects by augmenting the production of regulatory IL-10 in TLR 4 stimulated cells, which reduces inflammation and improves bacterial clearance in the spleen and not in the lungs [6]. Our data suggests the effects of binge alcohol may not only be dose dependent, but also tissue specific [46]. The tissue specific effects of binge alcohol and the modulating effects of cytokines are novel and interesting future studies that remain to be elucidated. Unlike binge alcohol dose administered mice infected with *B. thailandensis*, the spleen of mice infected with *B. vietnamiensis* did not express elevated IL-10 at 24 h PI. A mild anti-inflammatory response suggests differences in the host-pathogen interaction with *B. vietnamiensis* in which there is a greater tissue colonization and dissemination.

Although *B. vietnamiensis* is phylogenetically more distant to *B. pseudomallei* than *B. thailandensis*, it shares virulence factors with *B. pseudomallei* [49]. Extracellular lipase, metalloproteases, and proteases are thought to play roles directly related to the invasion and interaction with epithelial cells [31]. Relatedly, type I and type II secretion systems in *B. vietnamiensis* isolates were shown to be responsible for the secretion of proteins with hemolytic activity [18]. Similar to cystic fibrosis patients, pulmonary manifestations such as defective mucociliary function in binge drunk mice and humans predisposes to respiratory infections [25]. Our data illustrate that the evolutionary mechanisms utilized by *B. vietnamiensis* to colonize cystic fibrosis patients can be applied to our model of binge alcohol intoxication. Reinforcing these observations, Conway et al. (2004) have also shown that exopolysaccharide (EPS) produced by *B. cenocepacia* species interfered with phagocytosis of bacteria by human neutrophils and facilitated bacterial persistence in a mouse model of infection [12]. EPS was also found to inhibit neutrophil chemotaxis and production of oxygen reactive species [10]. Our mouse model intimates that binge alcohol creates a microenvironment in the host that could exacerbate infection by *Burkholderia* strains that are not commonly found in humans [1, 27].

In our study, *Burkholderia* near-neighbors were administered via the airways. We consider this route of infection more clinically relevant than subcutaneous or intraperitoneal injection. Nevertheless, humans usually acquire melioidosis by inoculation through skin abrasions, inhalation/aspiration and ingestion [48]. Although it is well documented that *B. pseudomallei* can be readily distinguished from *B. thailandensis*, the mechanism whereby the highly virulent *B. pseudomallei* causes disease in humans and animals is not well understood. Tan et al. (2008) speculated that a more lethal inhalational route of infection such as aerosolization is retained in the lungs, whereas an inoculum of bacteria delivered intranasally tends to be lodged in the nasal passage [48]. Our data indicate that binge alcohol consumption increases bacterial localization to the lungs and dissemination to the brain via an intranasal route of infection by 72 h PI. Neurological melioidosis is less frequently reported. *B. pseudomallei* can enter the brain and spinal cord via nasal branches of the trigeminal or olfactory nerve. Two alternative routes by which bacteria can reach the brain are via epithelial cell invasion and crossing the blood-brain barrier (BBB) [47]. To our knowledge, our study is the first to show *Burkholderia* near-neighbor distribution in to the brain. Our data suggest a singular binge alcohol episode could modulate nasal mucosa and related upper respiratory defenses that lead to greater pulmonary and neurological infections [47].

Interestingly, bacteria at low numbers were still observed in the lung and brain tissues of mice not administered alcohol and infected with *B. thailandensis* or *B. vietnamiensis*. These observations lead us to develop an *in vitro* system to test the effects of binge alcohol on lung and brain permeability and epithelial cell invasion. Epithelial and endothelial barrier integrity has a significant role in preventing bacterial translocation into the brain or other tissues [52]. To determine if a single binge alcohol dose increases lung and brain tissue permeability, we evaluated barrier leakiness using Evans blue dye 2 h post binge alcohol *in vivo* and discovered that the permeability of these tissues was induced by the alcohol (Figure 4). Consistent with this result, hazardous alcohol consumption in mice induces disruption of the colonic mucosal barrier that leads to leakage of bacterial toxins [34]. Likewise, primary cultured brain endothelial cells treated with 0.23 % (50mM) alcohol impairs BBB integrity *in vitro* [32]. In the present study, the TEER and FITC dextran flux rate were utilized to estimate the effects of binge alcohol on the paracellular permeability of lung epithelial (EpH4) and brain endothelial cell (bEndo.3) monolayers. The flow of ions in the paracellular gap was quantified by the TEER values. Our results showed that binge alcohol induced a significant decrease in TEER that remained suppressed for 8 h after a single alcohol treatment. A significant correlation was present between the decrease in the TEER value and the increase in the FITC dextran flux rate 2 h post binge alcohol administration. The increases in the FITC flux rate is especially relevant because infected mice treated with alcohol demonstrated elevated whole blood CFUs 2 h PI. It is possible that binge alcohol directly influences bacterial passage through epithelial or endothelial tight junctions [3]. However, it is unclear whether the impact of binge alcohol on tissue colonization is caused by modifications to epithelial raft structures, allowing for greater attachment, invasion, and bacterial survival; or synergy between transcellular and paracellular bacterial diffusion, aggravated by tight junction dysfunction [4].

Intracellular invasion of host cells is a successful survival strategy of many Gram-negative bacteria [29]. Furthermore, our previous data showed that low level alcohol exposure of *B. thailandensis* resulted in a reduction in motility and greater biofilm formation [26]. We speculated that these changes induced by alcohol could promote greater cellular attachment, invasion, and survival. Although *B. thailandensis* and *B. vietnamiensis* have been found intracellularly in both phagocytic and non-phagocytic cells, a quantitative study of alcohol-induced invasion of non-phagocytic epithelial cells has never been conducted. The results obtained in the present study demonstrated that binge alcohol significantly increased the invasion of both lung and brain non-phagocytic cells. Remarkably, less virulent *B. thailandensis* invaded lung epithelial cells at similar levels as *B. vietnamiensis* when cell monolayers were treated with alcohol during infection. Brain endothelial cells were more susceptible to *B. vietnamiensis* when exposed to alcohol. Our future studies will examine the effect of binge alcohol on the molecular mechanism of *Burkholderia* virulence and intracellular invasion with genomic and proteomics approaches [44, 54].

The data from the present study provide an important framework for *Burkholderia* near-neighbor virulence when patients engage in hazardous alcohol consumption. Our results showed that under binge alcohol conditions, intranasal infection with a less-pathogenic *B. thailandensis* can increase its infectivity, while diffusing into the blood stream. When using *B. vietnamiensis*, our findings indicate that a single binge alcohol episode increased *B. vietnamiensis* infectivity to a greater extent compared to *B. thailandensis*. Moreover, our findings provide novel insights into a possible mode of action for bacterial tissue colonization and dissemination via bacterial movement through paracellular space and intracellular invasion of non-phagocytic cells during binge alcohol exposure. The data from these studies support the public health responses being developed in melioidosis-endemic regions that emphasize the nature of alcohol consumption as a prime concern [14]. Emphasis is being placed on the dangers of binge drinking, especially around potential times of exposure to environmental *B. pseudomallei*, such as occurs with severe weather events and with certain occupations.

## Acknowledgements

We thank Northern Arizona University’s animal facility veterinarian and staff for providing guidance and training. We thank research associates at The Pathogen and Microbiome Institute and The Monroy lab at NAU for guidance and research support. In addition, we give our thanks to Dr. Nathan Nieto, from the Biology Department at NAU for helpful suggestions.

